# Heritability of hierarchical structural brain network

**DOI:** 10.1101/209635

**Authors:** Moo K. Chung, Zhan Luo, Nagesh Adluru, Andrew L. Alexander, Davidson J. Richard, H. Hill Goldsmith

## Abstract

We present a new structural brain network parcellation scheme that can subdivide existing parcellations into smaller subregions in a hierarchically nested fashion. The hierarchical parcellation was used to build multilayer convolutional structural brain networks that preserve topology across different network scales. As an application, we applied the method to diffusion weighted imaging study of 111 twin pairs. The genetic contribution of the whole brain structural connectivity was determined. We showed that the overall heritability is consistent across different network scales.

## 1. INTRODUCTION

In the usual brain connectivity studies, the whole brain is often parcellated into *p* disjoint regions, where *p* is usually 116 or less [1, 2]. For instance, Anatomical Automatic Labeling (AAL) parcellation provides 116 labels for all the cortical and subcortical structures (Figure 1) [1]. Subsequently, either functional or structural information is overlaid on top of AAL and *p* × *p* connectivity matrices that measure the strength of connectivity between brain regions are obtained. The major shortcoming of using the existing parcellations including AAL is the lack of refined spatial resolution. Even if we detected connectivity differences between large chunk of brain regions, it is not possible to localize what parts of parcellations are affected without additional analysis. There is a strong need to develop a higher resolution parcellation scheme.

**Fig. 1.**
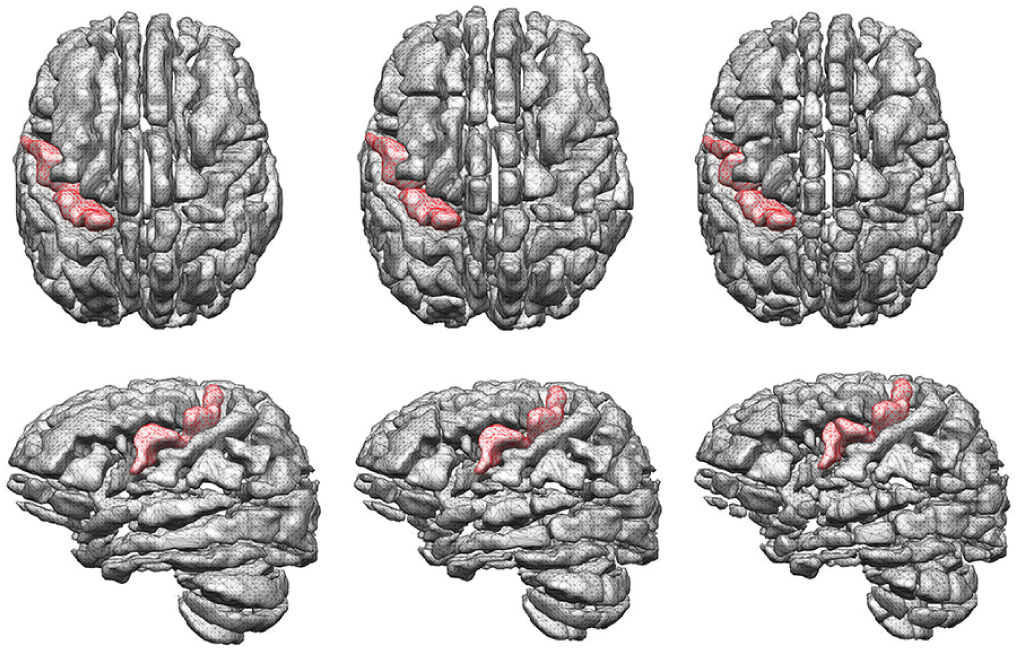
Left: AAL parcellation with 116 regions. Red region is the left precentral gyrus. Middle: the second layer of the hierarchical parcellation with 2 × 116 regions. Each AAL parcellation is subdivided into two disjoint regions. Right: the third layer of the hierarchical parcellation with 4 × 116 regions. Artificial gaps between subparcellations are introduced for visualization purpose only.

Brain networks are fundamentally multiscale. Intuitive and palatable biological hypothesis is that brain networks are organized into *hierarchies* [3]. A brain network at any particular sale might be subdivided into subnetworks, which can be further subdivided into smaller subnetworks in an iterative fashion. Unfortunately, many parcellation schemes give raise to conflicting topological structures of the parcellation from one scale to the next. The topological structure of parcellation at one particular scale may not carry over to different scales [2, 3]. There is a need to develop a hierarchical parcellation scheme that provide a consistent network analysis results and interpretation regardless of the choice of scale.

In this study, we propose a new hierarchical parcellation scheme based on the Courant nodal domain theorem [4]. The proposed method is related to graph cuts [5] and spectral clustering [6, 7] based parcellation schemes previous used in parcellating the resting-state functional magnetic resonance imaging (fMRI). However, in all these studies, parcellations are not hierarchical or nested so they produce conflicting topology over different network scales. Unlike previous approaches, our approach provides hierarchical nestedness and, thus, preserves topology across different spatial resolutions.

As an application, the proposed method was applied to diffusion weighted imaging (DWI) study of 111 twin pairs in determining the statistical significance of the genetic contribution of the whole brain structural connectivity.

## 2. HIERARCHICAL CONNECTIVITY

### Courant nodal domain theorem

For Laplacian ∆ in a compact domain 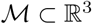, consider eigenvalues

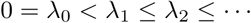

and eigenfunctions *ψ*_0_, *ψ*_1_, *ψ*_3_, … satisfying

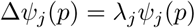

We then have 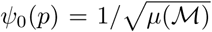, where 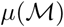 is the volume of 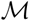. From the orthogonality of eigenfunctions, we have

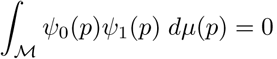

Thus, *ψ* _1_ must be take positive and negative values. The Courant nodal domain theorem [4] further states that *ψ*_1_ divides 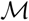 into two disjoint regions by the nodal surface boundary *ψ*_1_(*p*) = 0: When the domain is discretized as a 3D graph, the second eigenfunction *ψ*_1_ is called the Fiedler vector. It is often used in spectral clustering and graph cuts [5, 8]. Applying iteratively the nodal domain theorem, we can hierarchically partition 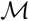 in a nested fashion.

### Hierarchical parcellation

The Courant nodal domain theorem is discretely applied to the AAL parcellation as follows. We first convert the binary volume of each parcellation in AAL into a 3D graph by taking each voxel as a node and connecting neighboring voxels. Using the 18-connected neighbor scheme, we connect two voxels only if they touch each other on their faces or edges. If voxels are only touching at their corner vertices, they are not considered as connected. This results in an adjacency matrix and the 3D graph Laplacian. The computed Fiedler vector is then used to partition each AAL parcellation into two disjoint regions (Figures 1 and 2). For each disjoint subregion, we further recompute the Fiedler vector restricted to the subregion. This binary partition process iteratively continues till all the partitions are voxels. We are doubling the number of parcellations at each iteration. There are a total of *p* = 116 parcellations in layer 1 and 2 ⋅ 115 parcellations in layer 2. At the *i*-th layer, there are 2^*i−*1^ ⋅ 116 parcellations. In our study, we were able to construct 20-layer nested hierarchical parcellations all the way to the voxel-level.

**Fig. 2.**
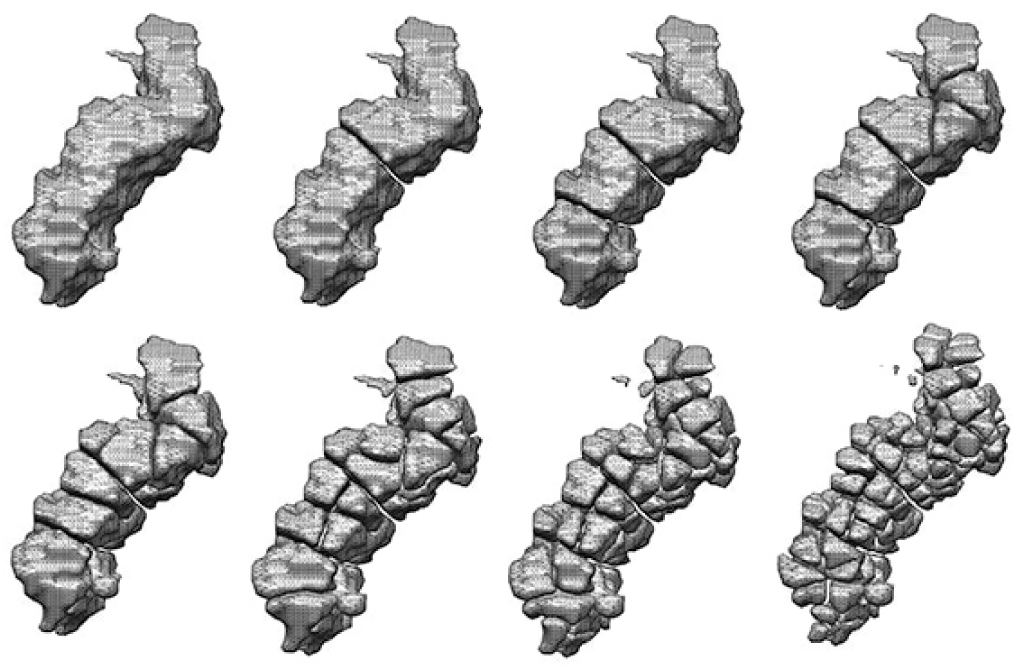
Hierarchical parcellation of the left precentral gyrus shown in Figure 1) up to the 8-th layer. At the 8-th layer, we have 2^8*−*1^ = 128 parcellations of the gyrus. The hierarchical parcellation continues till every voxel is a parcellation.

### Convolutional network

At the each layer of the hierarchical parcellation, we counted the total number of white matter fiber tracts connecting parcellations as a measure of connectivity. The resulting connectivity matrices form a *convolutional network*. Let 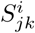 denote the total number of tracts between parcellations 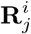 and 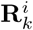 at the *i*-th layer. The connectivity 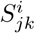 at the *i*-th layer is then the sum of connectivities at the (*i* + 1)-th layer (Figure 3), i.e.,

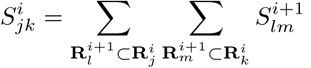

**Fig. 3.**
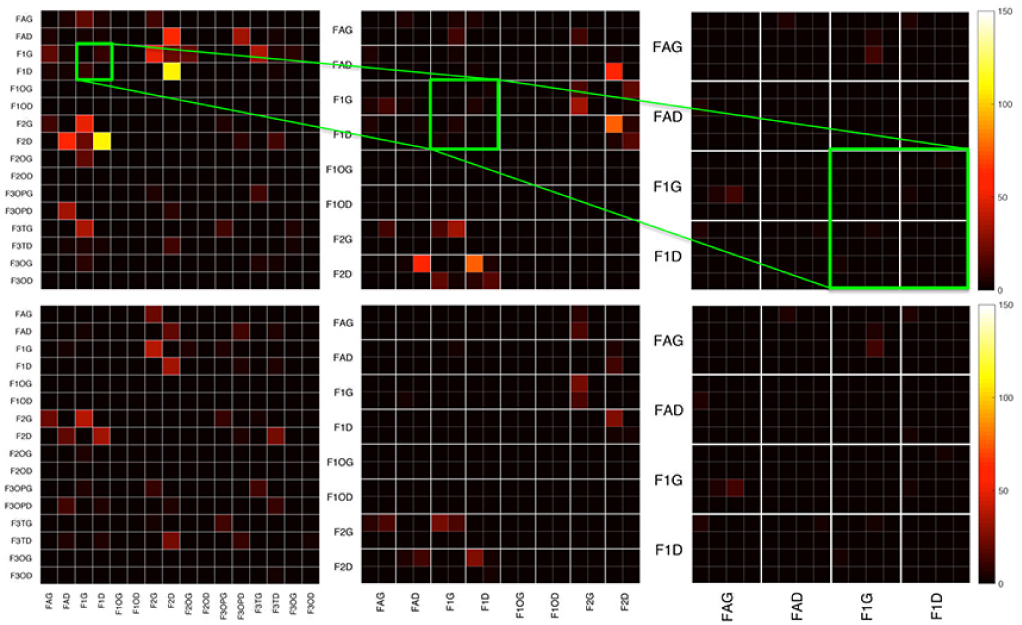
The hierarchical connectivity matrices of MZ-(top) and DZ-twins (bottom). The parts of connectivity matrices of the layers 1, 2 and 3 are shown. They form a layered convolutional network, where the convolution is defined as the sum of tracts between sub-parcellations.

The sum is taken over every subparcellation of 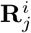 and 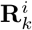

## 3. APPLICATION

### Subjects

Participants were part of the Wisconsin Twin Project [9]. 58 monozygotic (MZ) and 53 same-sex dizygotic (DZ) were used in the analysis. Twins were scanned in a 3.0 Tesla GE Discovery MR750 scanner with a 32-channel receive-only head coil. Diffusion tensor imaging was performed using a three-shell diffusion-weighted, spin-echo, echo-planar imaging sequence. A total of 6 non-DWI (b=0 s mm2) and 63 DWI with non-collinear diffusion encoding directions were collected at b=500, 800, 2000 (9, 18, 36 directions). Other parameters were TR/TE = 8575/76.6 ms; parallel imaging; flip angle = 90*◦*; isotropic 2mm resolution (128×128 matrix with 256 mm field-of-view).

Image preprocessing follows the pipeline established in [10]. FSL were used to correct for eddy current related distortions, head motion and field inhomogeneity. Estimation of the diffusion tensors at each voxel was performed using nonlinear tensor estimation in CAMINO. DTI-TK was used for constructing the study-specific template. Spatial normalization was performed tensor-based white matter alignment using a non-parametric diffeomorphic registration method. Each subject’s tractography was constructed using TEND algorithm, and tracts were terminated at FA-value less than 0.2 and deflection angle greater than 60 degree.

### Heritability Index

We are interested in knowing *the extent of the genetic influence on the structural brain network* of these participants and determining its statistical significance over different parcellation scales. For quantification, we used the heritability index (HI), which determines the amount of variation due to genetic influence in a population. HI is often estimated using Falconer’s formula as a baseline [11]:

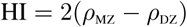

where *ρ*_MZ_ and *ρ*_DZ_ are the pairwise correlation between MZ-and and same-sex DZ-twins (Figure 4). For discrete tract counts, it is more reasonable to use Spearman’s correlation than Pearson’s correlation. The Pearson’s correlation does not work well with discrete tract count measures that often do not necessarily scale at the constant rate across different subjects and parcellations. Note Spearman’s correlation is Pearson’s correlation between the ordered tract counts.

**Fig. 4.**
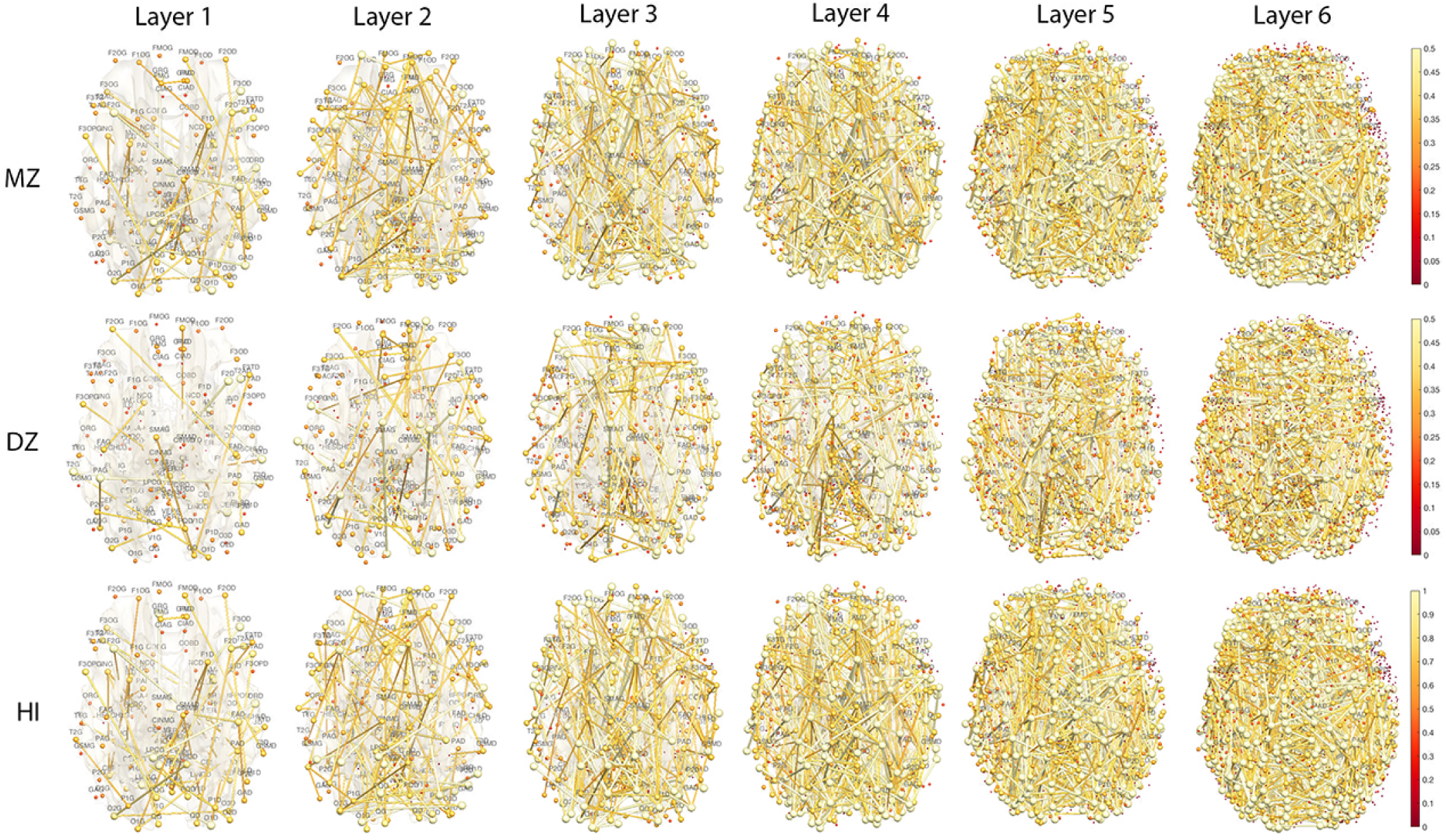
Top, middle: Edge colors are Spearman’s rank correlations thresholded at 0.3 for MZand DZ-twins for different layers. Node colors are the maximum correlation of all the connecting edges. Bottom: Edge colors are the heritability index (HI). Node colors are the maximum HI of all the connecting edges. MZ-twins show higher correlations compared to DZ-twins. The node and edge sizes are proportionally scaled.

### Exact Topological Inference

We determined the statistical significance of HI using the *exact topological inference* [11]. Consider weighted networks *G*^1^ and *G*^2^. Let 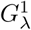 and 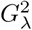 be the binary networks obtained by thresholding *G*^1^ and *G*^2^ at correlation *λ*. Let *B* be a monotonic graph function such that

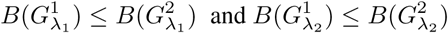

for *λ*_1_ ≤ *λ*_2_. The number of connected components (Betti-0 number) often used in persistent homology is such a function. The test statistic

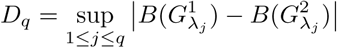

is used to determine the statistical significance. The thresholds *λ*_*j*_ are chosen uniformly in [0, 1] at 0.01 increment. The *p*-value under the null hypothesis of no network difference is then computed using [11]

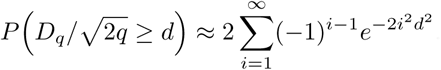

### Results

We are only showing results up to the 6-th layer, which has 3712 parcellations (61 voxels per parcellation in average). Beyond 6 layers, the individual parcellation was too small to be easily interpretable or visualize (Figure 4). At each layer, we performed the exact topological inference and obtained very consistent results. The *p*-values are less than 10^*−*12^ for each layer indicating the strong overall genetic contribution on the structural brain networks. Figure 5 shows the Betti-0 plots [11], which show the change of the number of connected components *B*(*G*_*λ*_) over correlation threshold values *λ* for MZ(solid yellow) and DZ-twins (dotted red). The sudden topological changes are occurring at the almost same correlation values regardless of the scale of the network.

**Fig. 5.**
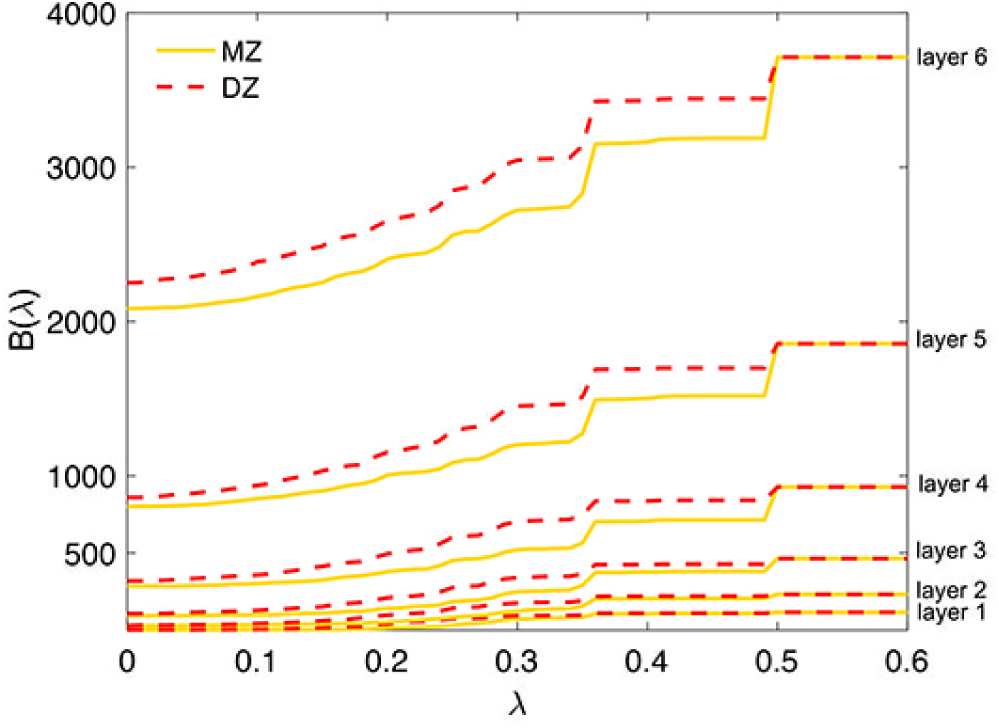
Betti-0 plots. The number of connected components (vertical) over the thresholded correlation values (horizontal) at each layer. The plots scale up over different layers resonably well. The sudden changes in the topological structure of network match up at the same correlation values.

## 4. CONCLUSION

We have developed new nested hierarchical structural brain parcellation and network methods. The methods were used in determining the genetic contribution of anatomical connectivity. The framework provides the topologically consistent statistical inference results regardless of the scale of the parcellation used. It is hoped the proposed parcellation and network construction frameworks will provide more consistent and robust network analysis across different studies and populations without concern for spatial resolution.

